# Tissue-specific Gene Expression Prediction Associates Vitiligo with SUOX through an Active Enhancer

**DOI:** 10.1101/337196

**Authors:** Zhihua Qi, Shiqi Xie, Rui Chen, Haji A. Aisa, Gary C. Hon, Yongtao Guan

## Abstract

Vitiligo is an autoimmune disease featuring destruction of melanocytes, which results in patchy depigemtation of skin and hair; two vitiligo GWAS studies identified multiple significant associations, including SNPs in 12q13.2 region. But one study ascribed the association to *IKZF4* because it encodes a regulator of T cell activation and is associated with two autoimmune diseases; while the other study ascribed the association to *PMEL* because it encodes melanocyte protein and has the strongest differential expression between vitiligo lesions and perilesional normal skins. Here we show that vitiligo associated gene in 12q13.2 region is *SUOX*. Reanalyzing one GWAS dataset, we predicted tissue-specific gene-expression by leveraging Genotype-Tissue Expression (GTEx) datasets, and performed association mapping between the predicted gene-expressions and vitiligo status. *SUOX* expression is significantly associated with vitiligo in both Nerve (tibia) and Skin (sun exposed) tissues. Epigenetic marks encompass the most significant eQTL of SUOX in both nerve and skin tissues suggest a putative enhancer 3Kb downstream of *SUOX*. We silenced the putative enhancer using the CRISPR interference system and observed 50% decrease in *SUOX* expression in K562 cells, a cell line that has similar DNase hypersensitive sites and gene expression pattern to the skin tissue at SUOX locus. Our work provided an example to make sense GWAS hits through examining factors that affect gene expression both computationally and experimentally.

## 1 Introduction

Gene expression is a desirable measurement in genetic association studies. First, according to NHGRI-EBI genome-wide association studies (GWAS) catalog, there are 46,159 SNPs identified through 4,201 GWAS in association with 2,364 traits (accessed on 21 February 2018), but majority of those GWAS SNPs are in non-coding region, and thus are believed to affect gene expression instead of protein function [1]. Second, many diseases manifest abnormal functions of specific tissue types [2], and consequently tissue-specific gene expression provides a better surrogate for genotypes in association mapping. Third, expression quantitative trait loci (eQTL), the genotype variants that affect gene expression, have been extensively studied [3, 4, 5], allowing gene expression to function as a bridge between genotypes and phenotypes. Unfortunately, the vast majority of existing GWAS datasets have no gene expression measurements at all, let alone tissue specific ones.

PrediXcan pioneered the idea of predicting gene expression [6], leveraging public resources such as the Genotype-Tissue Expression (GTEx) project [7], and performing gene-level association mapping with disease phenotypes. Bayesian variable selection regression (BVSR) excels at prediction, owing to its model averaging and shrinkage estimates [8]. Here we demonstrate that BVSR significantly outperforms Elastic-Net [9] (used by PrediXcan) in gene expression prediction. Our recent computational advancement makes BVSR applicable to large scale gene expression prediction [10]. BVSR provides the posterior inclusion probability (PIP) for each SNP, which measures the strength of marginal association in light of all SNPs, and PIP can be used to perform fine mapping.

We reanalysed a GWAS dataset of vitiligo, which is an autoimmune disease that features the destruction of skin pigment cells. Our analysis emphasize tissue-specific gene expression. We chose five tissue types as training dataset (Supplementary Figure S1(A)), and tissues are selected by jointly considering the number of samples available in GTEx, the origin of the germ layers, and their relevance to vitiligo. We predicted tissue-specific gene expression using BVSR, and performed gene-level association mapping between predicted gene expression and vitiligo status using logistic regression. For significant gene-level associations, we examined the predictive model to identify key predictor (SNPs with high PIP) to follow-up, including examining the methylation patterns and gene expression in cell lines after experimental interference. A highlight of our finding is the association between vitiligo and *SUOX*, whose expression is regulated by an nearby enhancer encompassing a significant GWAS hit.

## 2 Results

### Gene expression prediction

We trained BVSR models using GTEx datasets, predicted gene expression into GWAS datasets using the trained models, and performed association between predicted gene expression and phenotypes (details in Methods). To evaluate performance of BVSR in gene expression prediction, we used GTEx whole blood (*n* = 338) as training and Depression Genes and Networks (DGN, *n* = 922) as testing datasets. Samples in DGN have both genotypes and whole blood RNA-seq [5]; this allows us to compare the predicted against the measured gene expression using the coefficient of determination *R*^2^. Supplementary Figure S1(B) summarized the study overview.

To compare with Predixcan, we trained the model with Elastic-Net using the parameter settings described in their paper [7] for model fitting and cross-validation. Predixcan published all predication coefficients, but we have a different set of SNPs than what Predixcan used, which warrants the refitting. Figure 1 demonstrated a much improved performance of BVSR in gene expression prediction, compared with Elastic-Net. We attribute the improved performance to two technical aspects of BVSR. The first is model averaging. Intuitively, BVSR works harder to explore not only the best model, like Elastic-Net does, but also good models, and weights their contribution to prediction using the posterior probability of each model. The second is the separation of sparse and shrinkage priors. Such a separation allows sparse models without over-shrinking their parameter estimates. Three examples shown in Figure 1 appear to be the case where BVSR produced sparser models than Elastic-Net.

**Figure 1:**
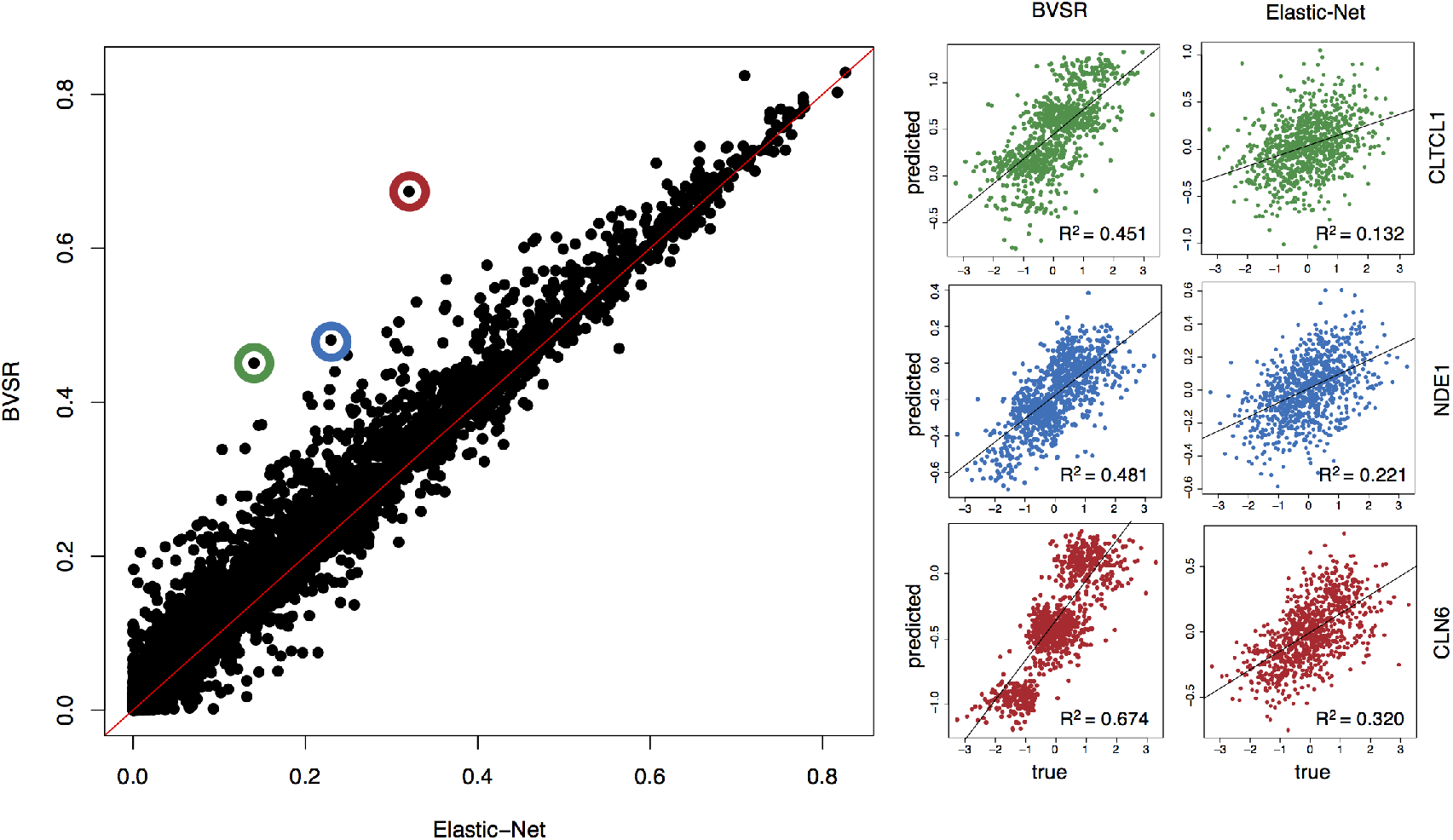
Comparison of predictive performance between BVSR and Elastic-Net. Using GTEx (n=338) as training and DGN (n=922) as testing, BVSR out-performs Elastic-Net in predicting gene expression. Three examples where BVSR outperforms Elastic-Net the most suggest BVSR has sparser predictive models.

### Vitiligo GWAS

Vitiligo is an autoimmune disease that features the destruction of melanocytes, resulting in patchy depigmentation of skin and hair [11]. It has an estimated worldwide prevalence of 1% [11], and the social or psychological distress caused by vitiligo can be devastating. We applied and downloaded 1, 251 vitiligo cases from dbGaP, and identified 4,155 healthy controls from other two GWAS datasets as controls for vitiligo (see Methods). Both cases and controls are European descents (Supplementary Figure S2). We first performed single SNP quality control separately for cases and controls, and after QC there are 514, 615 SNPs in cases and 495,103 in controls. Combined there are 523,349 SNPs, and these SNPs are subset of those GTEx SNPs. We used an in-house imputation software to fill in genotypes that are untyped in either cases or controls (See Methods). Poorly imputed SNPs were removed (Methods) and in the end we have 496, 847 SNPs for gene expression prediction and gene-level association mapping.

The gene-level association p-values appear to be well calibrated between any pair of five tissues (Figure 2). However, these p-values are inflated. The genomic control values range from 1.10 to 1.16 for each tissue, and the genomic control value is 1.13 for the combined p-values (Figure 2). We therefore adjust p-values using Benjamini-Hochberg-Yekutieli (BHY) procedure [12, 13, 14], separately for each tissue, to obtain corresponding q-values, each of which is the smallest FDR at which the hypothesis of interest would be rejected [15]. BHY procedure controls the false discovery rate assuming p-values are dependent, which is a welcoming feature for our gene expression prediction, as gene expressions are often correlated. Our simulations show that BHY procedure automatically corrects for p-value inflation featuring high genomic control values.

**Figure 2:**
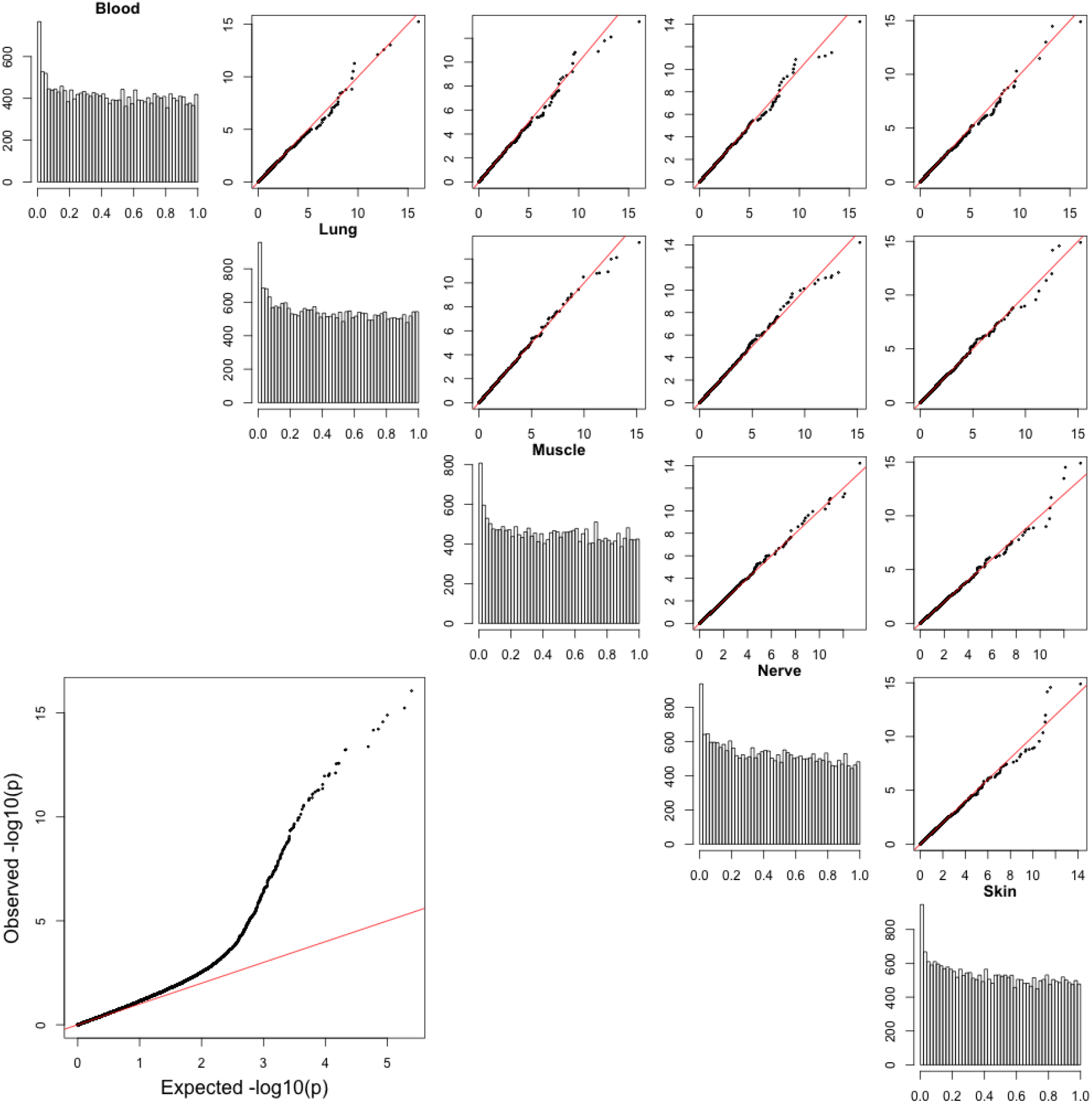
Calibration of association p-values. The diagonal plots are histograms for each set of p-values in five tissues. The upper off diagonal plots are pairwise quantile quantile plot of two sets of – log_10_(p-values). The lower off-diagonal plot is obtained by combining p-values from all five tissues. The lines y=x are colored in red.

At a nominal q-value cutoff of 0.05, our analysis discovered 6 genes outside of the MHC region whose predicted expression are significantly associated with vitiligo in at least two tissues (Table 1, Figure 3). Despite that vitiligo is a skin disease, the 6 genes appear to be lack of skin tissue-specificity: the nerve and skin tissues each has 5 associations, blood and lung each has 4, and muscle has 3.

**Figure 3:**
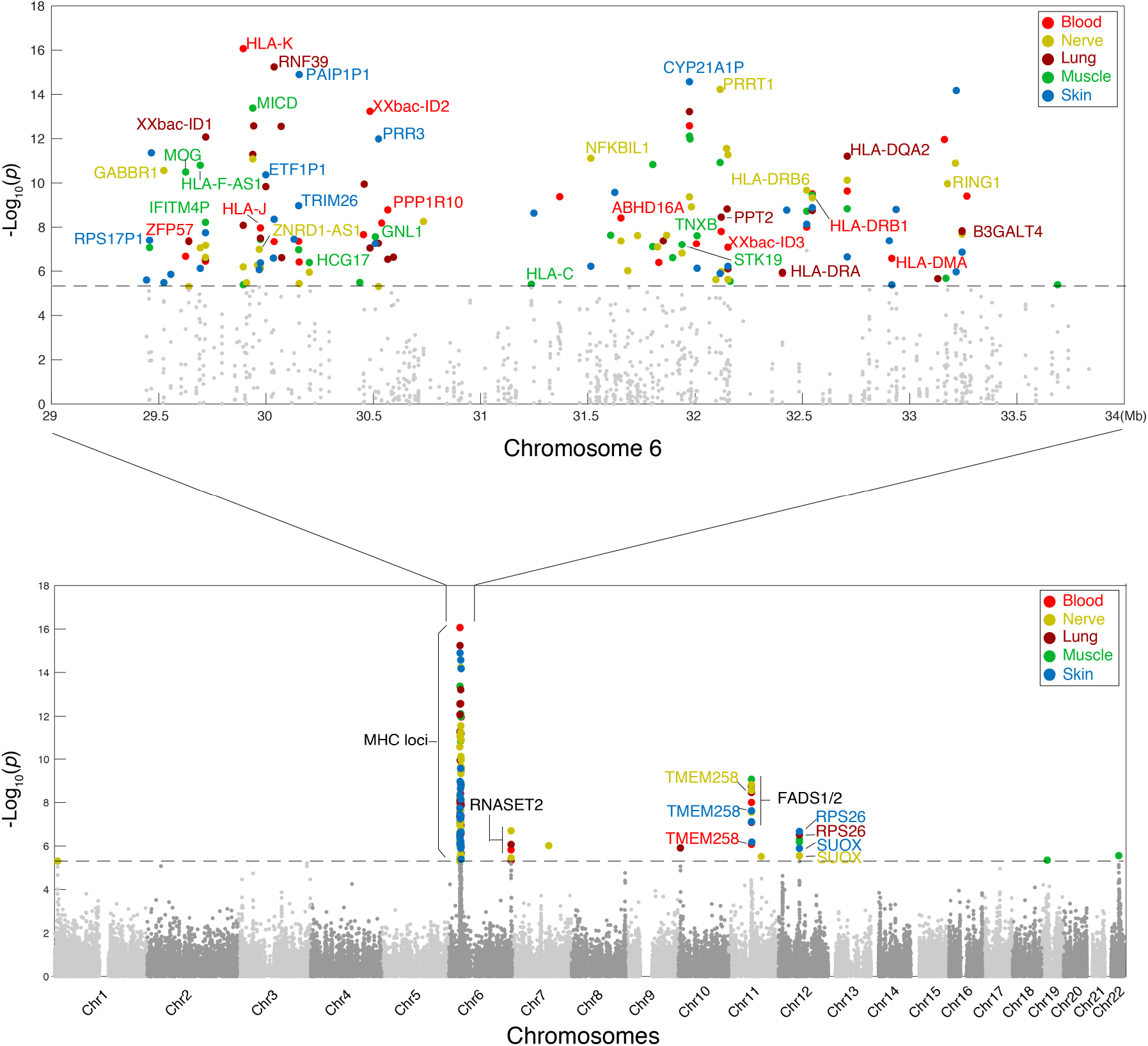
Manhattan plot of gene-level association in five different tissues for vitiligo. Different colors represent different tissues, whose GTEx datasets were used to train the predictive models. Y-axis is — log_10_p-value, although p-values are not used to call significance. Dashed horizontal line is the p-value cutoff of 5 × 10^−6^. Only genes that are significant (q-value < 0.05) in more than one tissue are labelled. At MHC region such a gene is only labelled once due to crowdedness. XXbac-ID1 stands for XXbac-BPG170G13.32, XXbac-ID2 for XXbac-BPG249D20.9, and XXbac-ID3 for XXbac-BPG300A18.13

**Table 1:** Vitiligo significant hits outside of the MHC region. P: p-values; Q: adjusted p-values using BHY procedure; OR: odds ratio. Position is the starting location of a gene and coordinates are from HG19. Significant test statistics (*Q* < 0.05) are marked bold. Only genes significant in more than one tissues are shown. Known GWAS hits are marked by †.

| Gene | Chr | Position | Statistics | Blood | Nerve | Lung | Muscle | Skin |
| --- | --- | --- | --- | --- | --- | --- | --- | --- |
| RNASET2 † | 6 | 167342992 | P | **1.4e-6** | **2.0e-7** | **8.5e-7** | 0.13 | **6.1e-6** |
|  |  |  | Q | **1.5e-2** | **2.5e-3** | **8.9e-3** | 1 | **4.4e-2** |
|  |  |  | OR | **0.83** | **0.82** | **0.83** | 0.94 | **0.84** |
| TMEM258 | 11 | 61535973 | P | **8.3e-7** | **2.8e-9** | **1.1e-6** | **5.7e-8** | **2.3e-8** |
|  |  |  | Q | **9.4e-3** | **6.1e-5** | **1.1e-2** | **1.0e-3** | **4.5e-4** |
|  |  |  | OR | **0.82** | **0.79** | **0.83** | **0.81** | **0.80** |
| FADS2 | 11 | 61560452 | P | **9.8e-9** | **2.7e-8** | **3.4e-9** | **8.5e-10** | **7.6e-8** |
|  |  |  | Q | **2.4e-4** | **4.7e-4** | **8.2e-5** | **3.0e-5** | **1.1e-3** |
|  |  |  | OR | **0.79** | **0.80** | **0.79** | **0.79** | **0.81** |
| FADS1 | 11 | 61567099 | P | **2.6e-8** | **1.4e-9** | 0.17 | **2.7e-9** | 0.10 |
|  |  |  | Q | **4.9e-4** | **3.4e-5** | 1 | **7.4e-5** | 1 |
|  |  |  | OR | **0.81** | **1.25** | 1.05 | **1.25** | 1.06 |
| SUOX | 12 | 56390964 | P | 6.2e-5 | **2.8e-6** | 8.2e-4 | 3.8e-3 | **1.2e-6** |
|  |  |  | Q | 0.34 | **2.4e-2** | 1 | 1 | **1.1e-2** |
|  |  |  | OR | 0.86 | **0.84** | 0.88 | 0.89 | **0.83** |
| RPS26 | 12 | 56435637 | P | 5.1e-6 | 3.2e-3 | **3.0e-7** | 0.36 | **2.1e-7** |
|  |  |  | Q | 5.1e-2 | 1 | **3.4e-3** | 1 | **2.9e-3** |
|  |  |  | OR | 1.18 | 1.11 | **1.20** | 1.03 | **1.21** |

Among the 6, *RNASET2* is known to be associated with vitiligo [16, 17]. The other 5 genes are clustered in two genomic regions. One region is in 11q12.2 that has three genes *TMEM258, FADS1*, and *FADS2. TMEM258* is required for full N-oligosaccharyl transferase catalytic activity in N-glycosylation. There is no immediate connection between its function to vitiligo, but glcosylation has a well known role in immune recognition. *FADS1* and *FADS2* are rate-limiting enzymes in the desaturation of linoleic acid to arachidonic acid, and alpha-linolenic acid to eicosapentaenoic acid and docosahexaenoic acid [18]. In humans, blood levels of polyunsaturated fatty acids (PUFAs) and long-chain PUFAs (LC-PUFAs) are strongly associated with *FADS1/2* [19]. Interestingly, PUFA of phospholipids reduction was observed in vitiligo epidermis [20], which provides a plausible link between *FADS1/2* and vitiligo. It is note-worthy that *FADS1* showed opposite effect sizes in blood tissue and never (or muscle) tissue.

The other region is in 12q13.2 that contains two genes: *SUOX* and *RPS26. RPS26* encodes a ribosomal protein that is a component of the 40S subunit; It is implicated in psoriasis, another skin disease marked by red, itchy, scaly patches [21]. *SUOX* encodes a enzyme, which is localized in mitochondria, that catalyzes the oxidation of sulfite to sulfate, the final reaction in the oxidative degradation of the sulfur amino acids cysteine and methionine. Melanogeneisis in cultured melanocytes can be substantially influenced by L-cysteine [22]. Sulfite induces oxidative stress, and in plants *SUOX* functions in sulfite detoxification and has been implicated in the adaption to elevated sulfur dioxide levels (e.g., acid rain) [23]. Epidermal melanocytes are particularly vulnerable to oxidative stress due to the pro-oxidant state generated during melanin synthesis [24].

Inside the MHC region our analysis discovered 36 genes at q-value cutoff of 0.05 (Figure 3). Among them only two genes, *HLA-DRA* and *HLA-DRB1*, were reported previously to be associated with vitiligo [25, 26]. There are 9 pseudogenes, including *IFITM?P, HLA-K*, and *MICD*, consisting 25% of the 36 genes, a proportion that is significantly higher than 16% pseudogenes in the MHC region and 11% in the whole genome. Associations inside the MHC region also appear not enriched in the skin tissue, which has 24 associations. As comparisons nerve has 23, lung 20, blood 19, and muscle 18. Interestingly, 9 genes show opposite effects in different tissues, including *GNL1, TNXB*, and *PPT2*. Supplementary Table S1 lists the 36 genes in the MHC region. A complete list of genes that are significant in at least one tissue can be found in the Supplementary Table S2.

### SUOX and an enhancer

*SUOX* is located in the 12q13.2 region identified by two vitiligo GWAS [27, 28]. The European study [27], however, ascribed the association signal to *IKZF4* because the most significant SNP rs1701704 is in its intron, and because *IKZF4* encodes a regulator of T cell activation, and *IKZF4* is associated with type 1 diabetes and alopecia areata. In the Chinese study [28], the most significant SNP is rs10876864, and the authors ascribed the association to *PMEL* based on two pieces of peripheral evidences: 1) *PMEL* encodes melanocyte protein and certain T cells exhibits reactivity to modified PMEL peptide epitopes in a subgroup of vitiligo patients; and 2) *PMEL* has the strongest differential expression between vitiligo lesions skin and vitiligo perilesional normal skin.

Our gene-level association ascribe the GWAS hits in 12q13.2 to *SUOX*. Figure 4 first presents a table detailing top 10 SNPs (with highest PIP) in the predictive models for the skin and nerve tissues. Very tellingly, seven overlapping SNPs in two predictive models are all eQTLs for *SUOX*, but only two SNPs are eQTLs for *PMEL* and none for *IKZF4*, according to GTEx portal (at FDR level of 0.05, accessed May 2018). The most significant eQTL in both skin and nerve — SNP rs10876864 — ranks first in predictive model for the nerve tissue and second for the skin tissue (Figure 4). Incidentally, this SNP is located in a region that is a DNase hypersensitive site in the skin tissue (Figure 5A). This led us to examine epigenetic marks in the cell line K562, which has similar DNase sensitivity in *SUOX* locus. Reassuringly, the gene expression patterns at the downstream of *SUOX* are also similar between K562 cells and the skin tissue. Moreover, both epigenetic markers H3K27ac and H3K4me3 have strong peaks near rs10876864, about 3kb downstream of the 3-UTR of *SUOX*, indicating this region to be a putative enhancer.

**Figure 4:**
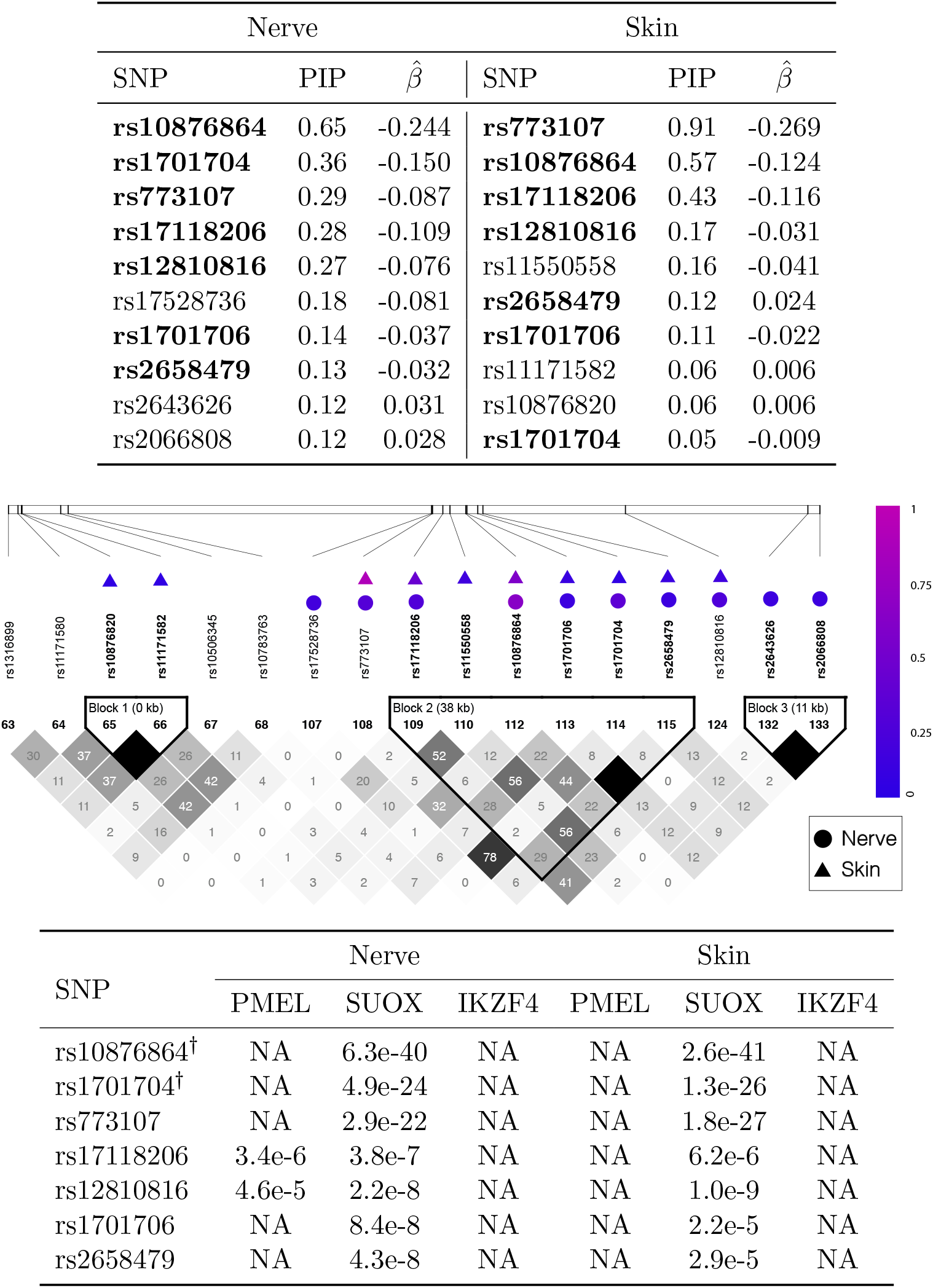
Fine mapping of SUOX in nerve and skin tissues. A) In each tissue, top 10 SNPs of highest PIP (posterior inclusion probability) are presented in the table, together with the predicting coefficients. (Both PIP and β are obtained from GTEx training dataset). Those 7 SNPs shared between two models (each for a tissue) are highlighted in bold. B) These top 10 SNPs are also marked in the LD plot, with the color of the dot representing the magnitude of PIP. The LD plot between SNPs was obtained from case control samples, where the number in each square represents correlation (×100) between two SNPs, ranging from 0 to 100 with 100 being darkest. C) The 7 shared SNPs are examined against GTEx portal for evidence of eQTLs in each tissue for three genes of interest, and p-values are provided if available. The entries marked by NA denote non-eQTL at FDR of 0.05. Two GWAS hit SNPs are marked by †.

**Figure 5:**
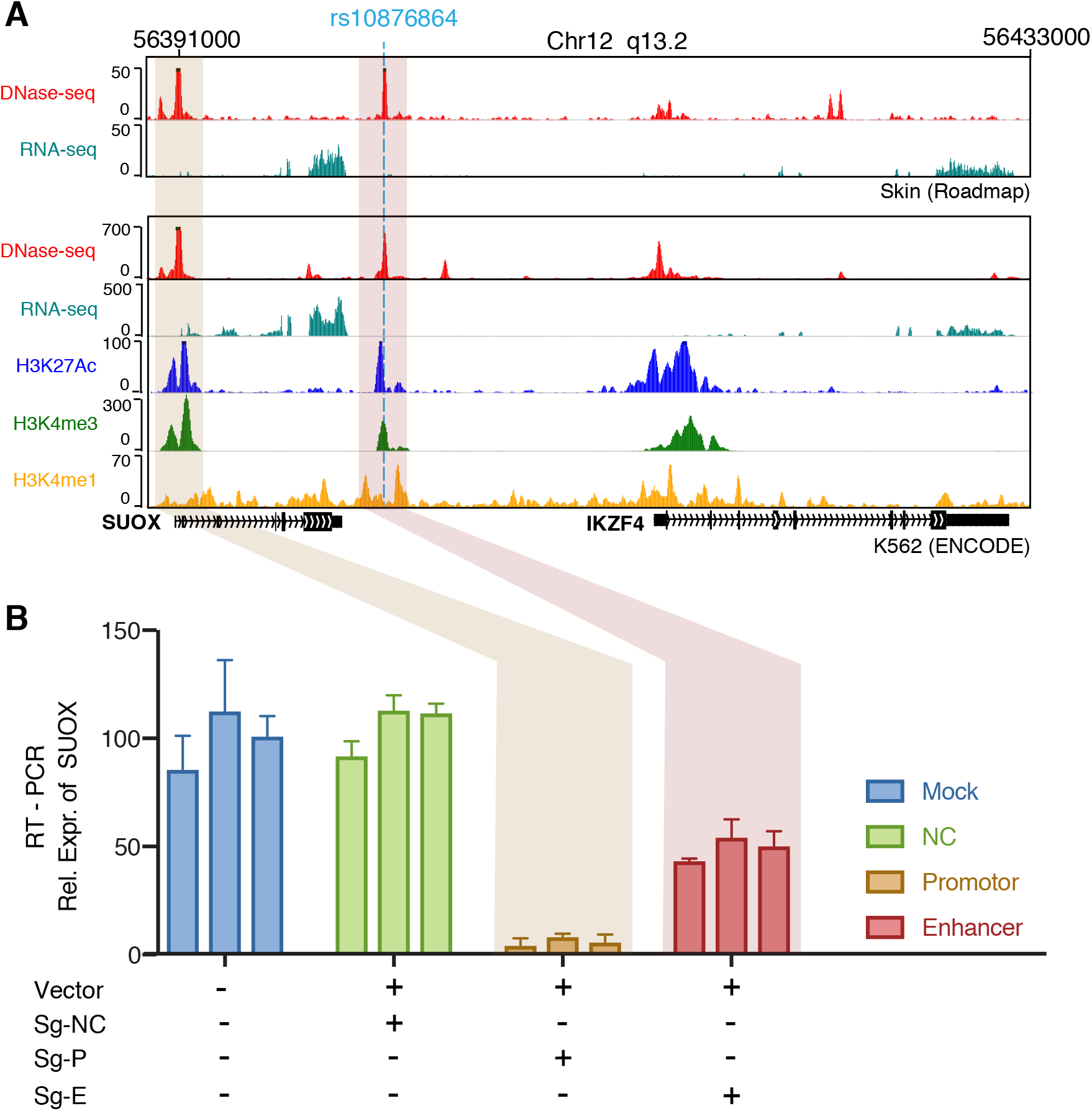
Epigenetic regulation of SUOX and its expression under CRISPR-i. A) Levels of transcription (mRNA) and DNase and epigenetic marks for the SUOX region in K562 cells and the Skin tissue. The position of SNP rs10876864 is marked by a blue dash vertical line. B) Using CRISPR-i to interfere SUOX promotor and an enhancer of interest, gene expression of SUOX were compared between mock (Vector), negative control (sg-NC), promotor-interfered (sg-P), and enhancer-interfered (sg-E).

We hypothesized that the putative enhancer affects gene expression of *SUOX*. To test the hypothesis, we used the CRISPR interference system[29] that can precisely silence a targeted promoter or enhancer. Taking advantage of an established CRISPR i system in K562 cell line [30], we designed two single guide RNA (sgRNA), one targets the promoter of *SUOX* as a positive control, and the other targets the putative enhancer. We also designed an sgRNA that targets *GFP* as a negative control, and performed mock experiment to measure *SUOX* expression as baseline for comparison. Figure 5B showed gene expression of *SUOX* under four experimental conditions. After the sgRNA that targets *GFP* was delivered with the CRISPR i, the gene expression measurement was similar to the baseline, suggesting negative control worked. When the sgRNA that targets the promoter was delivered with CRISPR i, the gene expression dropped to about 10% of the baseline level, suggesting that positive controls worked. Finally, when the sgRNA that targets the putative enhancer was delivered with CRISPR i, the gene expression dropped to about 50% of the baseline level, suggesting the putative enhancer did affect gene expression of *SUOX*.

## 3 Discussion

In this paper, we demonstrated the benefits of applying BVSR to predict tissue-specific gene expression and using predicted gene expression to perform association mapping. To make sense a significant SNP in GWAS, a common practice is to assign a gene based on the proximity to the SNP. This is apparently problematic. A famous example against such a practice is in obesity GWAS, which discovered an association SNP in the first intron of the *FTO* gene, but that SNP sits in an enhancer that promotes the expression of not *FTO* but *IRX3* that is 1Mbp downstream [31]. Most GWAS SNPs are intergenic or intronic and do not affect protein coding (or gene identity), rather, they associate with disease phenotypes through affecting gene expression [1]. Thus, focusing on gene expression to make sense GWAS hits — like we did here — is more advantageous. Our study provides a stellar example by showing that, in 12q13.2 region identified by two vitiligo GWAS, the relevant gene is not one of the ones reported by the GWAS studies (*IKZF4* and PMEL), but *SUOX*. Using CRISPR i system, we experimentally confirmed that an enhancer in 12q13.2 region, which is downstream of *SUOX*, mediates the expression of *SUOX*. The CRIPSR i system we used here appears to be ideal for functional assay of GWAS hits in noncoding region that are implicated in gene-level association.

SNP rs10876864 played a vital role in predicting gene expression of *SUOX*; it is also pivotal in locating the enhancer that mediates the expression of *SUOX*. In addition, SNP rs10876864 is a known GWAS hit of vitiligo [28]. Thus if it can be shown that different allele in SNP rs10876864 affects the potency of the enhancer, then we would close the remaining gap in elucidating the genetic association between rs10876864 and vitiligo. In fact, rs10876864 exhibited strong trans associations with 9 targets on 9 different chromosomes and in 4 distinct tissues: liver, omental adipose, blood cells and prefrontal cortex [32]. We hypothesize that the trans effects of rs10876864 is due to the very enhancer that mediate the gene expression of *SUOX*, which may also affect expression of other genes in trans.

## 4 Material and Methods

### 4.1 Datasets

We used the following datasets in our study.

- **GTEx:** We got tissue-specific RNA-seq and genome-wide genotype data from GTEx project (V6 release). We obtained five tissues from the dataset including whole blood (n= 338), nerve tibial (n=256), lung (n=278), skin (n=302) and muscle (n=361). The normalised gene expression was adjusted for sex, the top 3 principal components(PC) and the top 15 PEER factors (to quantify batch effects and experimental confounders)[7]. We used the GTEx dataset of different tissues to fit the predictive models.
- **DGN:** Depression Genetic Network (DGN) cohort obtained whole-blood RNA-seq and genome-wide genotype data for 922 individuals. For our analysis, we downloaded both the genotype and the HCP normalized RNA-seq data from the National Institute of Mental Health (NIMH) repository. DGN was used as a testing dataset in our study.
- **VitGene:** This vitiligo GWAS dataset contains only cases of 1251 samples (dbGaP accession number: phs000224). We obtained 4155 healthy controls from two GWAS datasets (dbGaP accession numbers: phs000336 and phs000147). The controls have the same continental origin with those cases (Supplementary Figure S2).

### 4.2 Genotype quality control and imputation

For each dataset, GTEx, DGN, case, and control, we performed single SNP QC separately. We excluded SNPs if the Hardy-Weinberg equilibrium exact test P-value < 1 × 10^−6^, or minor allele frequency, MAF < 1%. The cases and controls are typed on different platform. After QC there are 514, 615 SNPs in cases and 495,103 in controls. Combined there are 523, 349 SNPs, and these SNPs are subset of GTEx SNPs and HapMap3 SNPs.

In our experience, using popular imputation software, such as IMPUTE2 and MaCH, to combine datasets genotyped on different platforms is good for single SNP test, but often produces excessive false positives for multiple-SNP analysis or haplotype analysis. Thus, we performed imputation using our in-house software based on the two-layer model described in [33, 34]. Using HapMap3 as training datasets, our goal is to fill in the missing genotypes and to impute genotypes that are untyped in either cases or controls, but not both.

We first identified that there are 486, 396 SNPs shared between cases and controls, and masked those SNPs that are only typed in either cases or controls, then we thinned HapMap3 to contain only 523, 349 SNPs (the combined SNPs between cases and controls) and used these as training dataset to perform imputation to fill in the missing genotypes, including those that are only genotyped in either cases or controls. For those SNPs that are typed in one dataset but not the other, we can evaluate the imputation accuracy for that SNP, by comparing the genotyped and the imputed best guess genotypes. If the imputation error rate is > 5% this SNP will be removed. Otherwise, the SNP is kept. Where the imputed and the typed differ, we used typed. In the end, we have 496,847 SNPs for gene expression prediction.

### 4.3 Gene expression prediction using BVSR

For each gene, we used Bayesian variable selection regression to fit the following additive model ***Y** = **Xβ** + **ϵ** where **Y*** is an n-vector representing the individual gene expression, **X** is a *n × m* design matrix including m covariates (either SNPs or PCs), *β* is an m-vector and *ϵ* is the error term. After specifying sparse and shrinkage priors (detials can be found in [8]), we sampled different models to estimate β. And then applied this β to a design matrix of new set of (exchangeable) individuals to predict their gene expression. For each gene, we define *X* by including all SNPs 1 Mbp upstream of first exon and 1 Mbp downstream of the last exon. Note X is not allowed to have missing values; missing values are filled in by genotype imputation.

We used the software piMASS, the companions software of [8], to fit BVSR models. The software is run with the parameter of -w 10000, -s 1000000, -pmin 1, -pmax 10. After model fitting we use Rao-Blackwellized β estimates [8], weighted by the corresponding posterior inclusion probabilities, for prediction. For each gene, we fit the BVSR model twice to examine the uncertainty in prediction. We found highly congruent performance in all tissues except blood (Supplementary Figure S3). In real data analysis, for each gene we fit BVSR twice to obtain two predicted gene expressions, then we used the averaged gene expression to obtain a p-value for gene level association.

### 4.4 Gene expression prediction using Elastic Net

PrediXcan [6] used Elastic-Net to perform gene expression prediction. Elastic-Net linearly combines the *L*_1_ penalty (LASSO), with weight α and *L*_2_ penalty (ridge regression), with weight 1–*α*, to perform penalised regression [9]. Following the documentation in PrediXcan, we set *α* = 0.5 and applied 10-fold cross-validation to obtain λ, the combined penalty, to fit the penalised regression coefficients. The computation is done using *glmnet* package in R [35].

### 4.5 Predictive performance and gene-based association test

We used coefficients of determination (*R*^2^), between the predicted and measured values, to evaluate the predictive performance. For real data analysis, we predicted gene expression independently using each of the five GTEx tissues as training dataset. Because vitiligo has binary phenotypes, we fit a logistic regression between each predicted gene expression and disease phenotypes, controlling for sex and top ten PCs.

### 4.6 CRISPR interference experiments

#### 4.6.1 Cell Lines and Culture

K562 cells (from ATCC) were cultured in IMDM plus 10% FBS and pen/strep at 37 °C and 5% CO2.

#### 4.6.2 Plasmids

The lenti-sgRNA(MS2)-puro plasmid (Addgene ID: 73795) was used for sgRNA expression, and Lenti-dCas9-KRAB (Addgene ID: 89567) was used for dCas9-KRAB expression [30]. Two sgRNAs were designed for *SUOX* promotor (seq: GCCACCCGCTTCCAGCCAA; position: chr12:56391034-56391055) and putative enhancer (seq: ACGCCCGTAACGCAGC-CTC; position: chr12:56400869-56400888) respectively. The sgRNA fragments were inserted into the plasmid backbone (cut with BsmBl) by Golden Gate reaction. After transformation, single clone was picked and the sgRNA sequence of each clone was assessed by Sanger sequencing.

#### 4.6.3 Virus package

For lentiviral packaging, 3 × 10^6^ 293T cells were seeded in a 6cm dish one day before transfection. The indicated viral plasmid(s) were co-transfected with lentivirus packaging plasmids pMD2.G and psPAX2 (Addgene ID 12259 and 12260) with 4:2:3 ratios by using Lipofectamine 3000 (Thermo Fisher) according to the manufacturers protocol. Twelve hours after transfection, medium was changed to fresh DMEM with 10% FBS plus Pen/Strep. Seventy-two hours after transfection, virus-containing medium was collected, passed through a 45 *μ*m filter, and aliquoted into 1.5ml tubes.

#### 4.6.4 Virus titration

Viruses were stored in −80°*C* before infection or titration. The viruses are titrated by using the CellTiter-Glo luminescent cell viability assay. Briefly, 2 × 10^5^ cells were seeded into 96-well plate. Viruses are diluted in a 10-fold serial dilution and then used to infect the cells. The next day, medium was changed to fresh medium with antibiotics. Three days after infection, cells were harvested and the survival cell rate was identified by using CellTiter-Glo reagents. Based on the Poisson distribution, dilution with cell survival rate between 0% - 20% was used to back-calculate the virus titer.

#### 4.6.5 Virus infection

For virus infection, 2 × 10^5^ K562 cells were seeded in 24-well plate, and per well with 500 *μ*l complete medium plus 8*μ*g/ml polybrene. Virus aliquot(s) were thawed to room temperature and added to the plates. The plate was then centrifuged at 1000g at 36 ^°^C and returned to the incubator. The following day, the medium was changed to fresh complete medium with antibiotics to screen for infected cells. Cells were kept at 30% confluence during antibiotic selection.

#### 4.6.6 Quantifying SUOX expression

RNA was extracted using trizol (invitrogen)from K562 cells, infected by lentivirus with different targeting sgRNAs. Reverse transcription was performed using ReverTra Ace qPCR Master Mix(TOYOBO). SYBR green reagents were used for qPCR. Relavitve gene expression was analyzed using the 2-△△Ct method and normalized to that of *ACTIN* mRNA.

*SUOX* forward primer: GAAGACACTGGACCCGCAAAAG

*SUOX* reverse primer: GACTCGCAGGTGAACTCAGTG

*ACTIN* forward primer: GAGCACAGAGCCTCGCCTTT

*ACTIN* reverse primer: TCATCATCCATGGTGAGCTGG

## 5 Acknowledgement

Z.Q. is supported by a training fellowship from the Gulf Coast Consortia, on the NLM Training Program in Biomedical Informatics & Data Science (T15LM007093) NLM fellow. S.X. is an American Heart Association fellow (16POST29910007). H.A.A is supported by the Funds for the Xinjiang Key Research and Development Program (No. 2016B03038-1). G.C.H. is supported by the Cancer Prevention Research Institute of Texas (RR140023), the National Institutes of Health (DP2GM128203), the Department of Defense (PR172060), and the Welch Foundation (I-1926-20170325). Y.G is supported by United States Department of Agriculture/Agriculture Research Service under contract number 6250-51000-057 and National Institutes of Health under award number R01HG008157.

